# Nutrient availability-driven changes in extracellular matrix biochemical and mechanical properties regulate pancreatic cancer cell biology

**DOI:** 10.1101/2025.09.21.677637

**Authors:** Kristof Turan, Julien Guillard, Marta R. Storl-Desmond, Shan Lin, Yi-Chia Huang, Kevin Muñoz Forti, Simon Schwӧrer

## Abstract

The extracellular matrix (ECM) provides key biochemical and biomechanical cues that govern fundamental cellular processes, including growth and migration. ECM dysregulation and altered cell-matrix interactions are a driver of cancer progression, exemplified by pancreatic ductal adenocarcinoma (PDAC), where an abnormally dense, collagen-rich, and stiff ECM correlates with poor patient outcomes. The PDAC microenvironment is poorly perfused, resulting in altered nutrient availability, yet how this metabolic stress shapes the ECM and its biological activity remains poorly understood. Herein, we demonstrate that glutamine, a key amino acid depleted in poorly perfused PDAC regions, regulates the biochemical composition, mechanical properties, and biological activity of fibroblast-derived ECM. As glutamine availability increases, fibroblasts shift from producing a basement membrane-like ECM toward an interstitial, mature ECM enriched in fibrillar collagens. The ECM generated under glutamine-rich conditions is stiffer, which limits PDAC cell growth, while simultaneously, the elevated collagen I content promotes migration in a 3D spheroid model. Mechanistically, glutamine-dependent collagen I engages integrin α2 (ITGA2) to activate focal adhesion kinase signaling, driving PDAC cell migration independent of growth. In PDAC patients, glutamine stress inversely correlates with collagen expression in CAFs, with collagen I-ITGA2 as the most enriched ECM receptor interaction pair compared to the normal pancreas. These findings establish nutrient availability as a key regulator of ECM biology and offer new avenues to therapeutically intervene with dysregulated cell-matrix interactions in PDAC.

## Introduction

Pancreatic ductal adenocarcinoma (PDAC) is an aggressive tumor and is projected to become the second-leading cause of cancer-related mortality by 2030 in the United States [1,2]. PDAC is characterized by a prominent desmoplastic stroma, featuring aberrant amounts of extracellular matrix (ECM) [3]. Changes in ECM protein abundance and mechanical properties during tumorigenesis affect critical biological processes in cancer cells, such as their growth and migration [4]. Compared to a normal pancreas, the ECM in PDAC is enriched in collagens [5,6], especially collagens that are highly crosslinked, such as collagen I [7], resulting in an increased tumor stiffness [8,9]. Both increased stiffness and collagen I levels in PDAC tumors are associated with poor outcomes in patients [10–12]. Cancer cells sense ECM stiffness and collagen abundance through integrins and other cell surface receptors, and their engagement activates various mechanosensitive pathways, including focal adhesion kinase (FAK) [13]. Treatment of PDAC tumors with inhibitors against FAK proved effective in mouse models, delaying tumor progression [14]. These findings suggest targeting altered ECM signaling as a promising opportunity for cancer treatment.

Over 90% of PDAC ECM, and virtually all collagen, is produced by cancer-associated fibroblasts (CAFs) [7]. Despite the link between collagen I abundance and patient outcome, deletion of collagen I in CAFs promotes PDAC progression and worsens overall survival in mouse models [15], suggesting that CAF-derived collagen I restrains PDAC growth. In contrast, inhibition of collagen crosslinking and, thus, PDAC ECM stiffness reduces tumor progression in mouse models [9,12]. These findings indicate a disconnect between the effects of collagen I abundance and collagen-associated stiffness on PDAC cell behavior. Indeed, ECM dysregulation in cancer is now appreciated to play both pro-tumorigenic and anti-tumorigenic roles [4]. However, the cellular processes in PDAC differentially impacted by ECM dysregulation, and the precise ECM features that are tumor-promoting or tumor-restraining remain unclear.

Metabolism has emerged as a key regulator of fibroblast collagen synthesis [16]. The transition of fibroblasts towards CAF states is induced by cancer cell-derived factors, particularly transforming growth factor beta (TGFβ) [17]. It is accompanied by substantial metabolic reprogramming required for the efficient expression, translation, and stability of collagen [16]. For example, TGFβ-induced signaling promotes glutamine uptake, glutaminolysis, and utilization of the resulting glutamate for TCA cycle anaplerosis and proline biosynthesis, which are required for collagen I production [16,18]. Thus, TGFβ-induced collagen I synthesis in fibroblasts is glutamine-dependent [19,20]. In contrast, it is less well understood how glutamine availability impacts other matrisomal proteins, including other collagens, ECM glycoproteins, and proteoglycans. Despite the association of ECM stiffness with collagen I abundance and crosslinking, whether glutamine availability affects ECM mechanical properties is unknown. Glutamine availability contributes to intratumoral metabolic stress, with plasma levels of glutamine higher than in plasma than in PDAC interstitial fluid [21]. Furthermore, within PDAC, glutamine concentration varies spatially, forming areas with varied levels of metabolic stress [22]. How alterations in glutamine availability influence fibroblasts’ ability to produce a tumor-promoting or tumor-restraining ECM remains enigmatic.

Methods to produce fibroblast-derived ECM *in vitro* have been developed to model *in vivo*-relevant 3D cell-matrix interactions [23,24]. If produced from tissue-specific fibroblasts, these matrices are similar in composition to the tissue’s ECM from which they were isolated [25]. Moreover, ECM produced by CAFs can model key properties of tumor ECM, including elevated thickness, collagen content, stiffness, and anisotropy [26–28]. As a result, such ECMs evoke responses from cancer cells exposed to them that are reminiscent of the desmoplastic stroma [24]. For example, tumor-like ECM induces a spindle-shaped morphology, growth along aligned fibers, and promotes the migration of cancer cells [27–29]. These findings position fibroblast-derived ECN as a tool to better understand the differential effects of ECM composition and mechanical properties on PDAC cell biology. Despite these advances, tumor-relevant changes in nutrient availability have not yet been incorporated into fibroblast-derived ECM workflows, thereby limiting their *in vivo* relevance and, thus, their value for understanding ECM biology and cancer cell-matrix interactions. Here, we show that glutamine availability dictates the biochemical composition, mechanical properties, and biological activity of fibroblast-derived ECM, resulting in differential impacts on PDAC cell growth and migration.

## Results

To understand whether glutamine availability in PDAC is associated with differential expression of ECM components, we analyzed CAFs extracted from a publicly available single-cell RNA-sequencing (scRNA-seq) dataset of human PDAC patients [30] for the expression of core matrisome gene sets [31] and genes associated with glutamine stress in fibroblasts [19] (Sup. Fig.1A). Projecting both the fibroblast glutamine stress signature and matrisome signatures onto uniform manifold approximation and projection (UMAP) of all PDAC CAFs (Sup. Fig.1B) revealed an inverse relationship between glutamine stress and collagen expression, but not with ECM glycoproteins and proteoglycans (Sup. Fig. 1C-J). These data suggest that glutamine stress in fibroblasts may result in different ECM expression patterns. To directly test whether glutamine availability impacts the biochemical composition of fibroblast-derived ECM, we analyzed ECMs produced by murine pancreatic stellate cells (PSCs), CAF precursor cells in PDAC [32], in the presence of TGFβ under high (4 mM, HQ), medium (1 mM, MQ), or low (0.4 mM, LQ) glutamine concentrations via LC-MS-based proteomics (Fig. 1A). We chose this range because 0.4 mM glutamine is at the lower end of glutamine concentrations found in PDAC tumor interstitial fluid [21,33], and 4 mM is the glutamine concentration in DMEM, a standard culture medium. 120 matrisome proteins were detected, with contributions from both core matrisome and matrisome-associated categories (Fig. 1B). Principal component analysis separated ECMs by glutamine concentration (Fig. 1C). The largest source of variance in the dataset (PC1, explaining 47.9%) corresponded to differences between the high and medium/low glutamine concentrations, indicating that glutamine availability substantially affected matrisome proteins.

**Figure 1:**
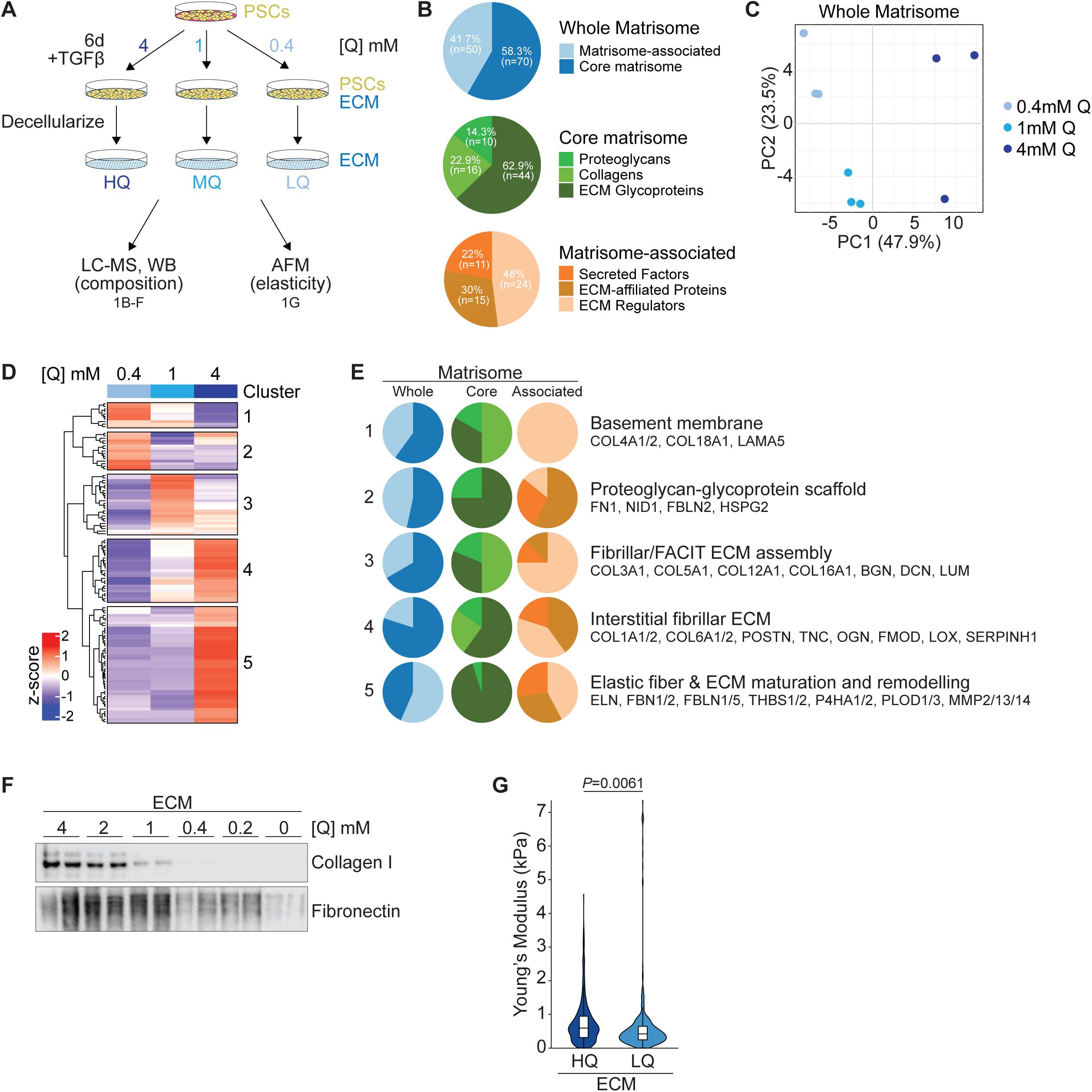
Glutamine availability during ECM synthesis regulates ECM composition and stiffness. (A) Schematic of fibroblast-derived ECM production under high (HQ; 4 mM), medium (MQ, 1 mM) or low (LQ; 0.4 mM) glutamine (Q) concentration in the presence of 2 ng/mL TGFβ for 6 days, followed by decellularization and the indicated downstream analyses. (B) Matrisome classification of ECM proteomics data. Pie charts summarize the proportion of matrisome categories detected by LC-MS. (C) Principal component analysis (PCA) of matrisome proteins in ECM proteomics data, derived from precursor intensities for matrisome proteins generated at 4-, 1-, and 0.4 mM glutamine (Q). *n*=3 biological replicates. (D) Heatmap of matrisome proteins in ECM proteomics data showing z-score average precursor intensities for matrisome proteins across 4-, 1-, and 0.4 mM glutamine (Q). Five hierarchical clusters are highlighted following the calculation of Euclidean distance. *n*=3 biological replicates. (E) Pie charts summarizing the proportion of matrisome categories across the five defined clusters (D) with a brief content summary and representative members. Cluster number is indicated on the left. *N*=10 proteins (cluster 1), *N*=15 proteins (cluster 2), *N*=24 proteins (cluster 3), *N*=25 proteins (cluster 4), *N*=46 proteins (cluster 5). (F) Western blot of decellularized fibroblast-derived ECM produced across glutamine titration (4, 2, 1, 0.4, 0.2, 0 mM). ECMs were collected in the same volume, and equal volumes unadjusted for protein concentrations were loaded. *n*=2 biological replicates. Representative images are shown. (G) Young’s modulus of ECMs produced at 4 mM (HQ) or 0.4 mM (LQ) glutamine, measured by AFM. Data show the distribution of individual indentations as violin plots; white boxes indicate median and interquartile range. *n*=448 measurements (HQ); *n*=528 measurements (LQ), derived from *n*=5 biological replicates per group. *P*-values were calculated by unpaired two-tailed *t* test.

Hierarchical clustering of matrisome proteins by Euclidean distance resolved five clusters with distinct matrisome contents (Fig. 1D, E; Sup. Fig. 2A). Two clusters were LQ-biased and captured basement-membrane proteins (Cluster 1) and associated proteoglycan/glycoprotein scaffolds (Cluster 2); notably, Cluster 2 does not comprise collagens. A third cluster peaked at the MQ condition and included fibrillar collagens and fibril-associated collagens with interrupted triple helices (FACIT) as well as several small leucine-rich proteoglycans (SLRPs), consistent with a transitional state in collagen fiber organization [34]. The remaining two clusters contained proteins that increased with glutamine concentration and were enriched for interstitial, fibrillar collagens (Cluster 4), elastic fiber constituents (Cluster 5), and ECM maturation enzymes (Cluster 4 and 5); strikingly, Cluster 5 also lacked collagens but comprised enzymes involved in fibrillar collagen hydroxylation and remodeling, consistent with ECM maturation. Protein-protein interaction network analysis of each cluster supported these classifications (Sup. Fig. 2A). Amino acid composition analysis of the proteins across clusters revealed differences in proline and glycine content (*P*<0.01, Sup. Fig. 2B), the major building blocks of collagens, consistent with the absence of collagens in Clusters 2 and 5 (Fig. 1E). Protein glutamine content was comparable across clusters (Sup. Fig. 2B), indicating roles for glutamine beyond its direct incorporation into ECM proteins.

With hierarchical clustering showing fibrillar collagens as highly glutamine-responsive (Sup. Fig. 2C) while a glycoprotein scaffold persisted at LQ (Fig. 1D, E), we chose collagen I and fibronectin, respectively, for validation studies. Of note, ECM samples analyzed by Western blot could not be adjusted for total protein content, which was 47.6% (0.4 mM) and 68.1% (1 mM) of that of ECM HQ (4 mM), according to LC-MS peptide determination (data not shown). Western blot of decellularized ECM produced under a glutamine titration confirmed that the reduction of glutamine from 4 mM to 1 mM was sufficient to reduce collagen I abundance, with no substantial effects on fibronectin (Fig. 1F). While fibronectin levels also decreased as the glutamine concentration was lowered, this effect was much less pronounced compared to collagen I, which was essentially depleted at 0.2 mM glutamine (Fig. 1F). Accounting for the differences total ECM protein supports the finding that fibronectin is relatively enriched in ECM LQ compared collagen I, which is enriched in ECM HQ. The same trend was observed in ECM derived from human PSCs (Sup. Fig. 2D), providing cross-species validation. The enrichment of interstitial fibrillar collagens and enzymes involved in collagen hydroxylation and crosslinking in ECM HQ-biased Clusters 4 and 5 (Fig. 1E) indicates that glutamine availability may also impact ECM stiffness and, thus, the ECM’s mechanical properties. Consistently, atomic force microscopy (AFM) nano-indentation demonstrated that ECM LQ was less stiff (more elastic) than ECM HQ (Fig. 1G).

The finding that glutamine availability alters the biochemical composition and mechanical properties of fibroblast-derived ECM suggests that it may also impact its biological activity. To test this idea, we plated EGFP-labeled murine PDAC cells or spheroids derived from the KPC (*Kras*^LSL-G12D/+^;*Trp53*^LSL-R172H/+^;*Pdx1*-Cre) mouse model [31] on decellularized ECM produced under HQ and LQ conditions (Fig. 2A). ECM HQ restrained the growth of KPC cells compared to 2D culture (Fig. 2B). This effect was not apparent on ECM LQ (Fig. 2B), indicating that the growth restraining properties of ECM HQ cannot solely be explained by its 3D nature. Next, KPC-derived spheroids were transferred onto the ECM, resulting in the spreading of peripheral cells and migration into the ECM [26] (Sup. Fig. 3). We used spheroid spreading on ECM HQ and LQ over time as a readout for migration. In contrast to growth, spheroid spreading was promoted by ECM HQ compared to ECM LQ (Fig. 2C), a phenotype particularly apparent within the first 24h after spheroid transfer.

**Figure 2:**
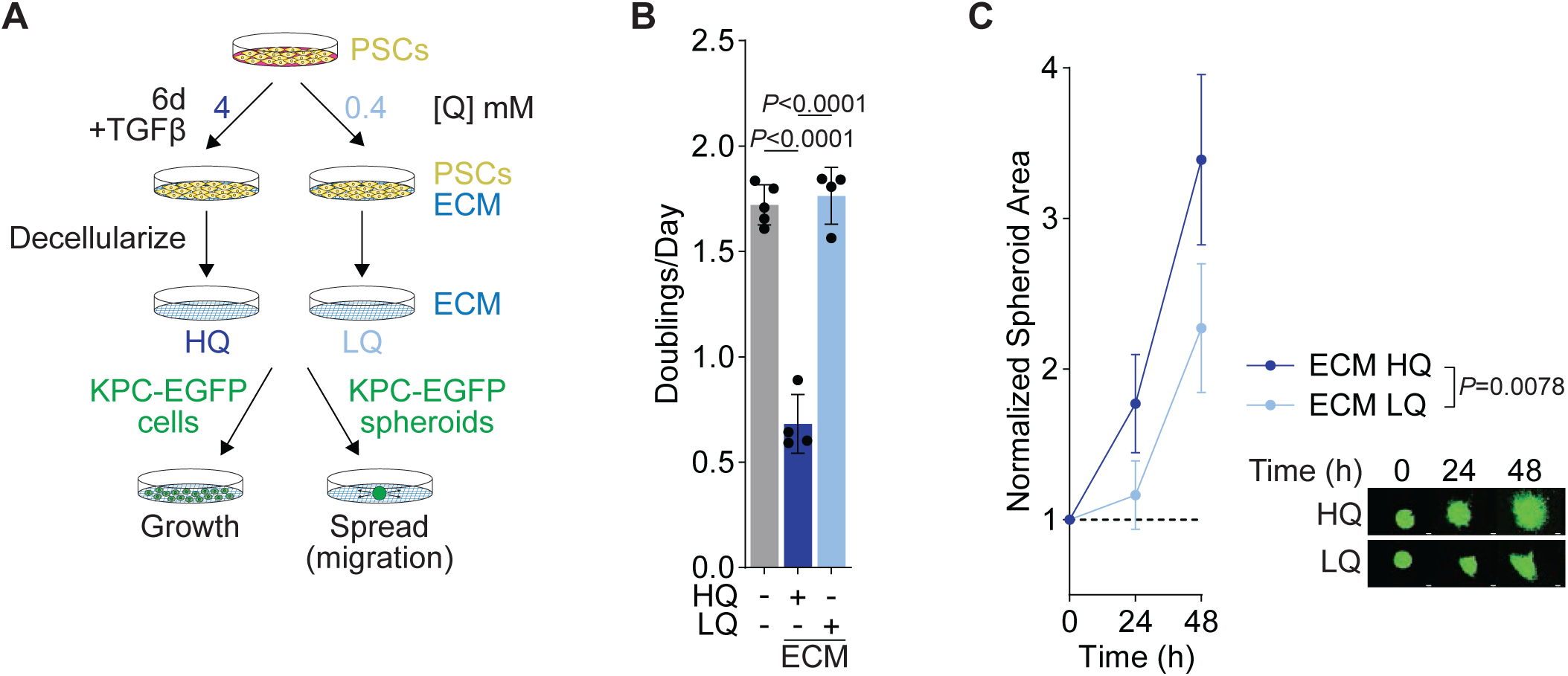
Glutamine availability during ECM synthesis alters the ECM’s biological activity. (A) Schematic of fibroblast-derived ECM production under 4 mM (HQ) and 0.4 mM (LQ) glutamine availability in the presence of 2 ng/mL TGFβ for 6 days, followed by decellularization and the indicated downstream assays to assess the ECM’s biological activity. (B) Growth of KPC2-EGFP cells on plastic, ECM HQ and ECM LQ over 48 hours. *n*=5 (Plastic), *n*=4 (ECM) biological replicates. Data represent mean ± SD. *P* values were calculated by one-way ANOVA. (C) Spreading of KPC2-EGFP spheroids on ECM HQ and ECM LQ over 48 hours. Each spheroid’s area at a given time point was normalized to its initial area after transfer to ECM. *n*=5 biological replicates. Data represent mean ± SD. *P* values were calculated by two-way ANOVA. Representative images of EGFP+ spheroids on ECM HQ and ECM LQ over time. Scale bar=100 µm.

We hypothesized that altered fibrillar collagen content and/or stiffness of ECM HQ and ECM LQ (Fig. 1) may underlie changes in PDAC cell behavior on these surfaces (Fig. 2). ECM stiffness is primarily driven by fibrillar collagen crosslinking mediated by lysyl oxidase (LOX) [9,35], which we found enriched in HQ-biased matrisome clusters (Fig. 1E), alongside fibrillar collagens. We reasoned that LOX inhibition during ECM HQ production would allow us to interrogate the role of ECM stiffness independent from its collagen content. To this end, we used PSCs to generate ECM under HQ in the presence or absence of β-aminopropionitrile (BAPN) (Fig. 3A), a natural and irreversible pan-LOX inhibitor [34]. BAPN treatment slightly increased levels of collagen I in the ECM (Sup. Fig. 4A). Notably, BAPN-treatment during ECM synthesis essentially rescued the growth restraining properties of ECM HQ (Fig. 3B) but did not inhibit KPC spheroid spreading on ECM HQ (Fig. 3C). To interrogate the role of collagen I content on PDAC cell behavior, we produced a non-cell ECM by creating a 3D gel containing varying concentrations of collagen I supplemented with a constant amount of Matrigel to provide additional ECM cues (Fig. 3D). Non-cell ECM with high collagen I content reduced the growth of KPC cells compared to ECM with low collagen I content or 2D culture (Fig. 3E), albeit to a lower extent than ECM HQ. Consistent with the observation that collagen I-rich ECM HQ drives migration, high collagen I levels promoted spheroid spreading on non-cell ECM (Fig. 3F). Collagen I coating of culture dishes was sufficient to reduce KPC growth and to promote KPC spheroid spreading (Sup. Fig. 4B-D), indicating that a 3D environment is not required for collagen I to exert its effects on PDAC cells.

**Figure 3:**
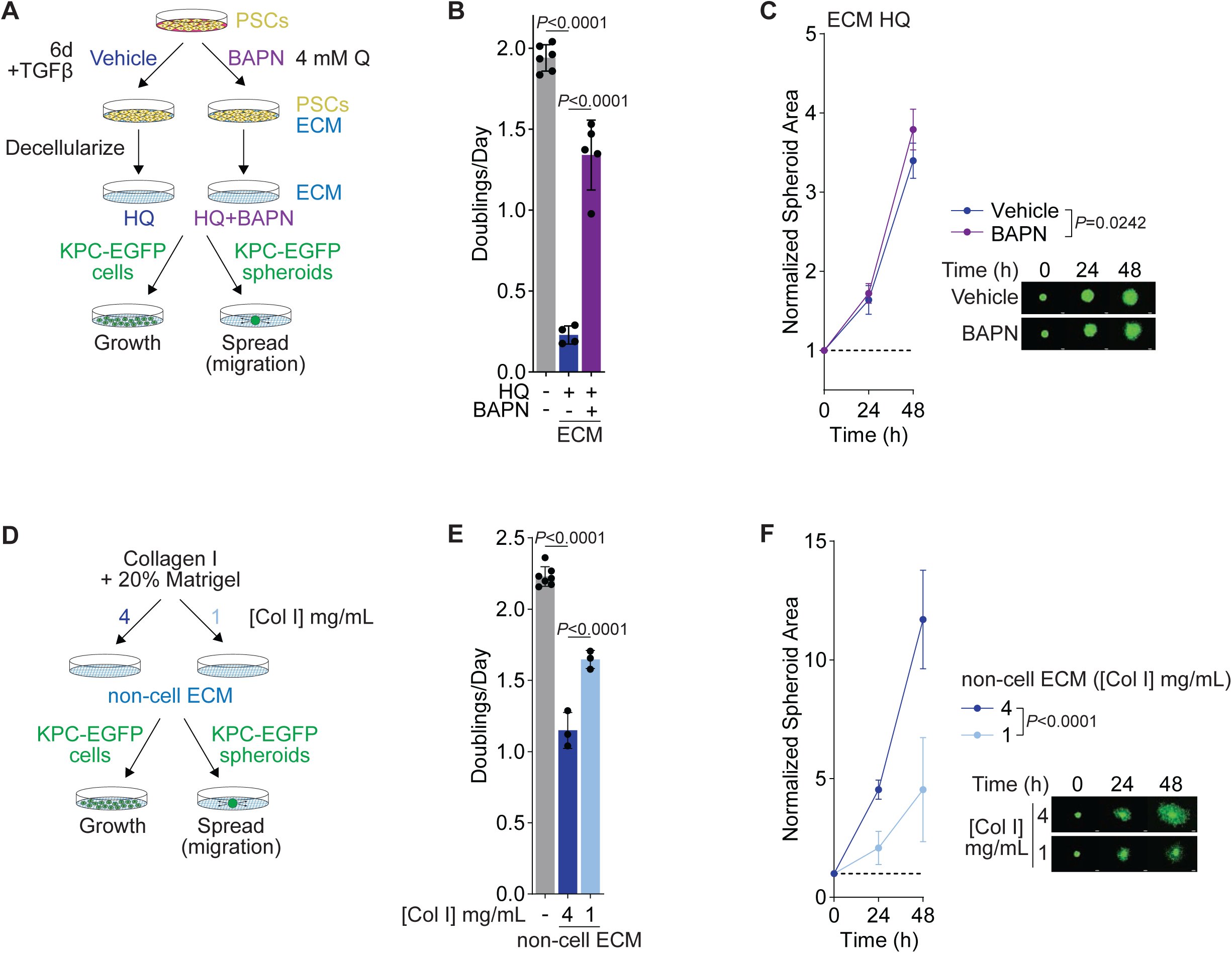
Collagen crosslinking and abundance underlie ECM HQ’s growth-restraining and pro-migration roles. (A) Schematic of ECM production under high glutamine (HQ; 4 mM) in the presence of 2 ng/mL TGFβ for 6 days, accompanied by either vehicle or 250 µM β-aminopropionitrile (BAPN) treatment to inhibit LOX-mediated collagen crosslinking. ECMs were then decellularized and used for the indicated assays. (B) Growth of KPC2-EGFP cells on ECM HQ prepared with vehicle or BAPN treatment (plastic control included). *n*=6 (plastic), *n*=4 (ECM HQ, vehicle), *n*=5 (ECM HQ, BAPN) biological replicates. Bars show mean ± SD. *P* values were calculated by one-way ANOVA. (C) Spreading of KPC2-EGFP spheroids on ECM HQ prepared with vehicle or BAPN treatment over 48 hours. Spheroid area at each time point was normalized to its own initial area after transfer to ECM. *n*=6 biological replicates. Data represent mean ± SD. *P* value was calculated by two-way ANOVA. Representative images of EGFP+ spheroids on ECM HQ vehicle and ECM HQ BAPN over time. Scale bar=100 µm. (D) Schematic of synthetic ECM assays: 3D gels composed of collagen I at two concentrations (4 mg/mL or 1 mg/mL) were mixed with 20% Matrigel and then seeded with KPC2-EGFP cells for growth or spheroids for spreading assays. (E) Growth of KPC2-EGFP cells on plastic or synthetic ECM prepared with 4 or 1 mg/mL collagen I. *n*=7 (plastic), *n*=3 (synthetic ECM) biological replicates. Bars represent mean ± SD. *P* values were calculated by one-way ANOVA. (F) Spreading of KPC2-EGFP spheroids on synthetic ECM prepared with 4 or 1 mg/mL collagen I over 48 hours. Areas were normalized to each spheroid’s initial area (0 h). *n*=6 (4 mg/mL), *n*=5 (1 mg/mL). Data represent mean ± SD; *P* value was calculated by two-way ANOVA. Right: representative images of EGFP+ spheroids on synthetic ECM with 4 or 1 mg/mL collagen I over time. Scale bar=100 µm.

We turned to the Pancreatic Ductal Adenocarcinoma study (PAAD) from The Cancer Genome Atlas (TCGA) to determine potential pathways underlying the migration-promoting effects of collagen I-rich ECM HQ. We ranked patients based on their combined expression of *COL1A1* and *COL1A2*. Gene set enrichment analysis (GSEA) revealed an enrichment of FAK and MAPK signaling in patients with high *COL1A1/2* expression (Fig. 4A). Consistently, culture of KPC cells on collagen I-rich ECM HQ but not collagen I-poor ECM LQ induced FAK and MAPK signaling (Fig. 4B). MAPK signaling on ECM HQ was sensitive to FAK inhibition (Fig. 4C), indicating that ECM HQ-induced FAK signaling underlies MAPK signaling in KPC cells. The two FAK inhibitors PF-562271 and PND-1186 impaired KPC spheroid spreading on ECM HQ (Fig. 4D) but did not alter KPC growth on ECM HQ (Fig. 4E). Consistent with FAK being upstream of MAPK signaling, the MEK inhibitor Trametinib also inhibited KPC spheroid spreading but only slightly reduced KPC growth on ECM HQ (Fig. 4D, E).

**Figure 4:**
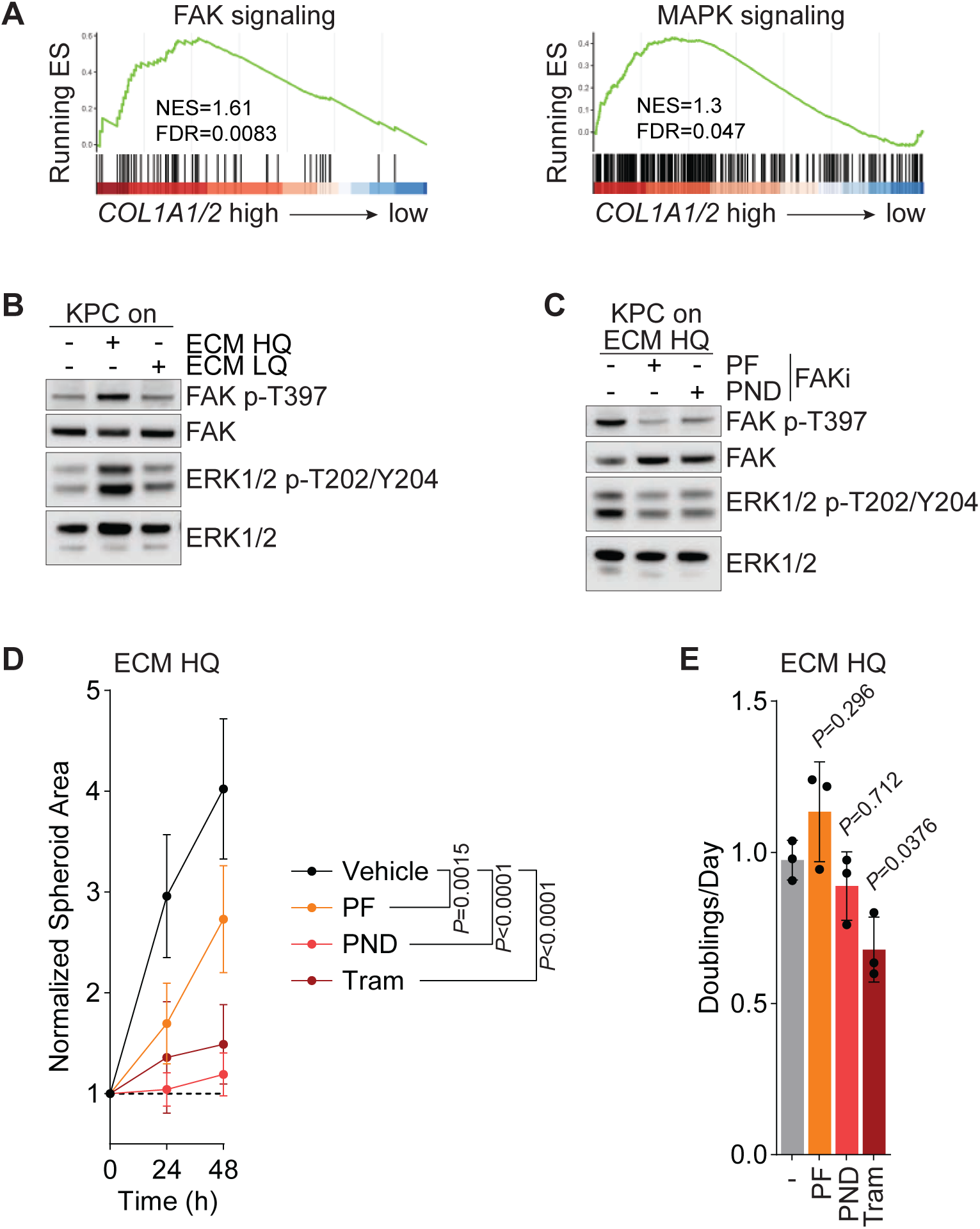
FAK/MAPK signaling mediates PDAC cell migration downstream of ECM HQ. (A) Gene set enrichment analysis (GSEA) of patients (tumors) from The Cancer Genome Atlas (TCGA) Pancreatic Ductal Adenocarcinoma (PAAD) study ranked by mean *COL1A1*/2 expression. (B) Western blot for FAK and MAPK signaling in KPC2-EGFP cells 24h after plating on plastic, ECM HQ, or ECM LQ. Representative image. (C) Western blot for FAK and MAPK signaling in KPC2-EGFP cells 24h after plating on ECM HQ in the presence of two different FAK inhibitors (FAKi) - 1 µM PF-573228 (PF) or 1 µM PND-1186 (PND). Representative image. (D) KPC2-EGFP spheroid spreading on ECM HQ in the presence of FAK or MEK inhibitors. Normalized spheroid area over 48 h for vehicle, PF (1 μM), PND (1 μM) and Trametinib (0.1 μM) treatment. Each spheroid’s area at a given time point was normalized to its own initial area after transfer to ECM. *n*=6 biological replicates. Data represent mean ± SD. *P* values were calculated by two-way ANOVA. (E) Growth of KPC2-EGFP cells on ECM HQ over 48 h for vehicle, PF (1 μM), PND (1 μM) and Trametinib (0.1 μM) treatment. *n*=3 biological replicates. Bars show mean ± SD. *P* values were calculated by one-way ANOVA.

The above data suggest that ECM HQ promotes PDAC cell migration by activating FAK signaling. We hypothesized that collagen I, which is enriched in ECM HQ and sufficient to promote PDAC cell migration (Fig. 1E, F, Fig. 3F, Sup. Fig. 4D), is sensed by cell surface receptors on PDAC cells to activate FAK and induce migration. To identify relevant collagen receptors, we performed CellChat analysis [36] to infer cell-matrix interactions from human PDAC scRNA-seq data [30] (Sup. Fig. 5A). This analysis revealed that interactions with tumor epithelial cells are the strongest among the fibroblast interactome (Sup. Fig. 5B). Furthermore, fibroblasts had the highest probability to communicate with tumor epithelial cells through collagens across all cell types (Fig. 5A). Such interactions included several fibroblast-derived collagens and different receptors on tumor epithelial cells, including various integrins and syndecans (Fig. 5B). Among these, predicated collagen interactions with the *ITGA2/ITGB1* and *ITGA3/ITGAB1* heterodimers were specific to tumor epithelial cells (Sup. Fig. 5C). We also leveraged a dataset comparing changes in putative cell interactions between the normal pancreas and pancreatic tumors [37]. This analysis revealed *COL1A1*-*ITGA2* and *COL1A2*-*ITGA2* as the most upregulated ligand-receptor pair between fibroblast collagens and epithelial cells in PDAC tumors compared to the normal pancreas (Fig. 5C). Consistently, *COL1A1* and *COL1A2* were highly expressed by fibroblasts (Sup. Fig. 5D), and *ITGA2* expression was restricted to tumor epithelial cells (Sup. Fig. 5E). *ITGA2* expression correlated with *COL1A1/2* expression and was associated with poor outcomes in TCGA-PAAD (Fig. 5D, E). To test whether ITGA2 is required for ECM HQ to activate FAK/MAPK signaling, we generated KPC cells with *Itga2* deletion. *Itga2* knockout reduced FAK and MAPK signaling in KPC cells on ECM HQ and collagen I-coated plates (Fig. 5F, G), which was rescued by expression of inducible human *ITGA2* in KPC cells with *Itga2* deletion (Fig. 5H).

**Figure 5:**
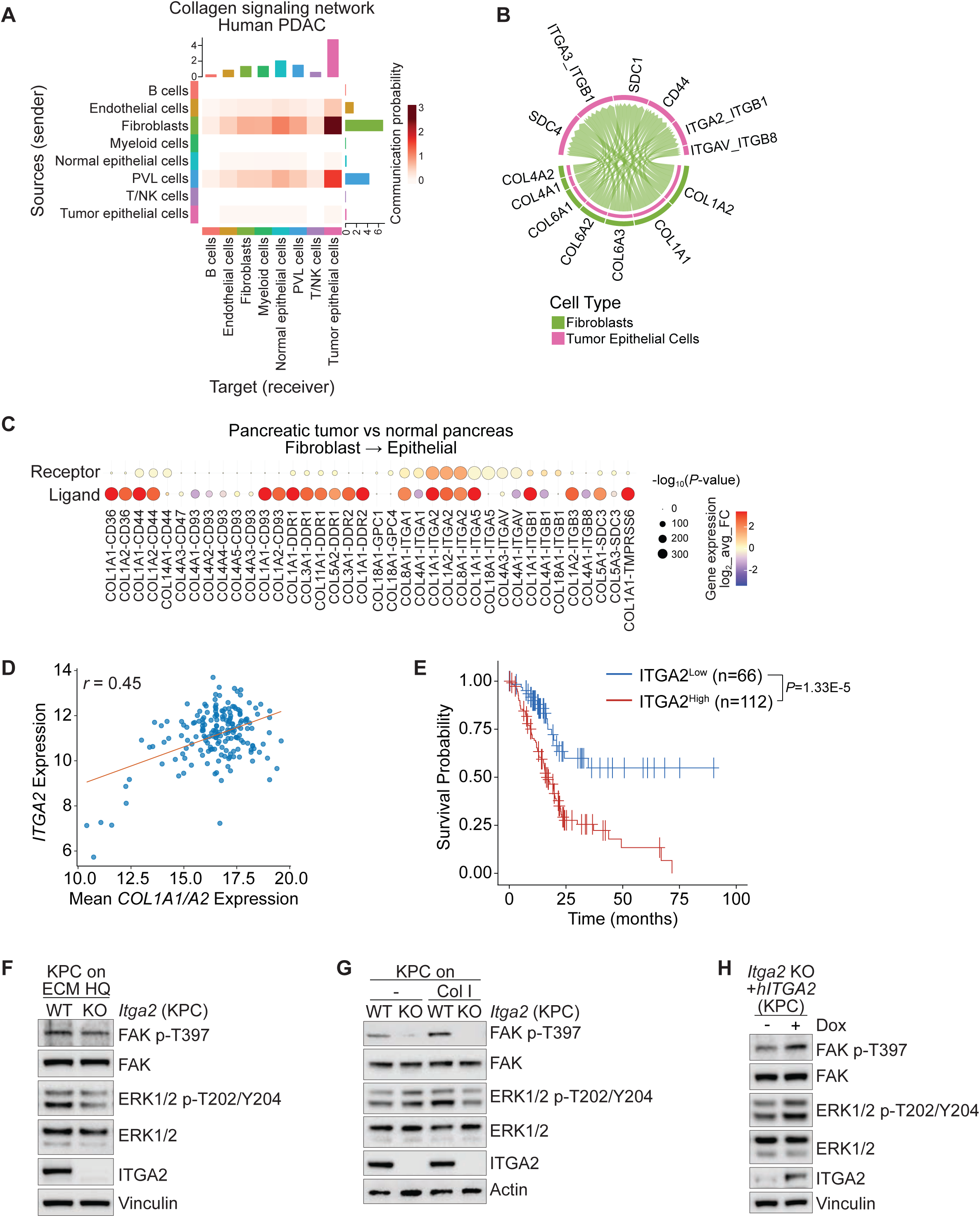
Collagen I-ITGA2 interactions are enriched in PDAC and promote FAK/MAPK signaling in PDAC cells on ECM HQ. (A) Collagen signaling network inferred from human PDAC scRNA-seq (PRJCA001063). Heatmaps show source (sender) and target (receiver) communication probabilities by cell type. (B) Chord diagram of collagen ligand–receptor pairs for predicted fibroblasts (green) to tumor epithelial cells (pink) signaling from (A). (C) Bubble plot showing differential collagen signaling in tumor versus normal pancreas for the fibroblast→epithelial route (from [37]). Average expression of respective ligands (fibroblast) and receptors (epithelial cell) across all cells from the same type is shown. (D) Pearson correlation between *ITGA2* and mean *COL1A1/2* expression in TCGA PAAD bulk RNA-seq (one dot per tumor). (E) Overall survival of patients in TCGA PAAD stratified by *ITGA2* expression. Sample split was determined by the optimal cut-off method. (F) Western blot of KPC2-EYFP cells wildtype (WT) or knockout (KO) for *Itga2*, plated on ECM HQ for 24h. Representative image. (G) Western blot of KPC2-EYFP *Itga2* WT or KO cells, 24h after plating on plastic or collagen I-coated plates. Representative image. (H) Western Blot of KPC2-EYFP *Itga2* KO cells expressing doxycycline (DOX)-inducible *hITGA2*, in the presence or absence of 1 µg/mL DOX, on ECM HQ for 24h. Representative image.

With ITGA2 regulating migration-promoting FAK/MAPK signaling in response to fibroblast-derived ECM, we next sought to understand the role of ITGA2 in PDAC cell behavior on ECM HQ. *Itga2* deletion abrogated KPC spheroid spreading on ECM HQ and non-cell ECM with high collagen I content (Fig. 6A, B). While *Itga2* deletion also reduced spheroid spreading on ECM LQ, this effect was much less pronounced compared to ECM HQ (Sup. Fig. 6A). Similarly, *Itga2* was required for spheroid spreading in 2D only in the presence of collagen I (Fig. 6C), consistent with a surface-specific role. This effect was rescued by expression of human *ITGA2* (Sup. Fig. 6B, C). While *Itga2* deletion reduced KPC cell growth on collagen I-coated dishes, it had no impact on growth on ECM HQ (Fig. 6D). Consistent with the genetic data, inhibition of ITGA2-ITGB1 interaction with BTT-3033 reduced spheroid spreading on ECM HQ in a dose-dependent fashion (Fig. 6E), while slightly but significantly promoting KPC cell growth on ECM HQ (Fig. 6F). Finally, we used a neutralizing antibody to block ECM interactions with ITGA2 (also called CD49b), which was sufficient to reduce spheroid spreading on ECM HQ (Fig. 6G).

**Figure 6:**
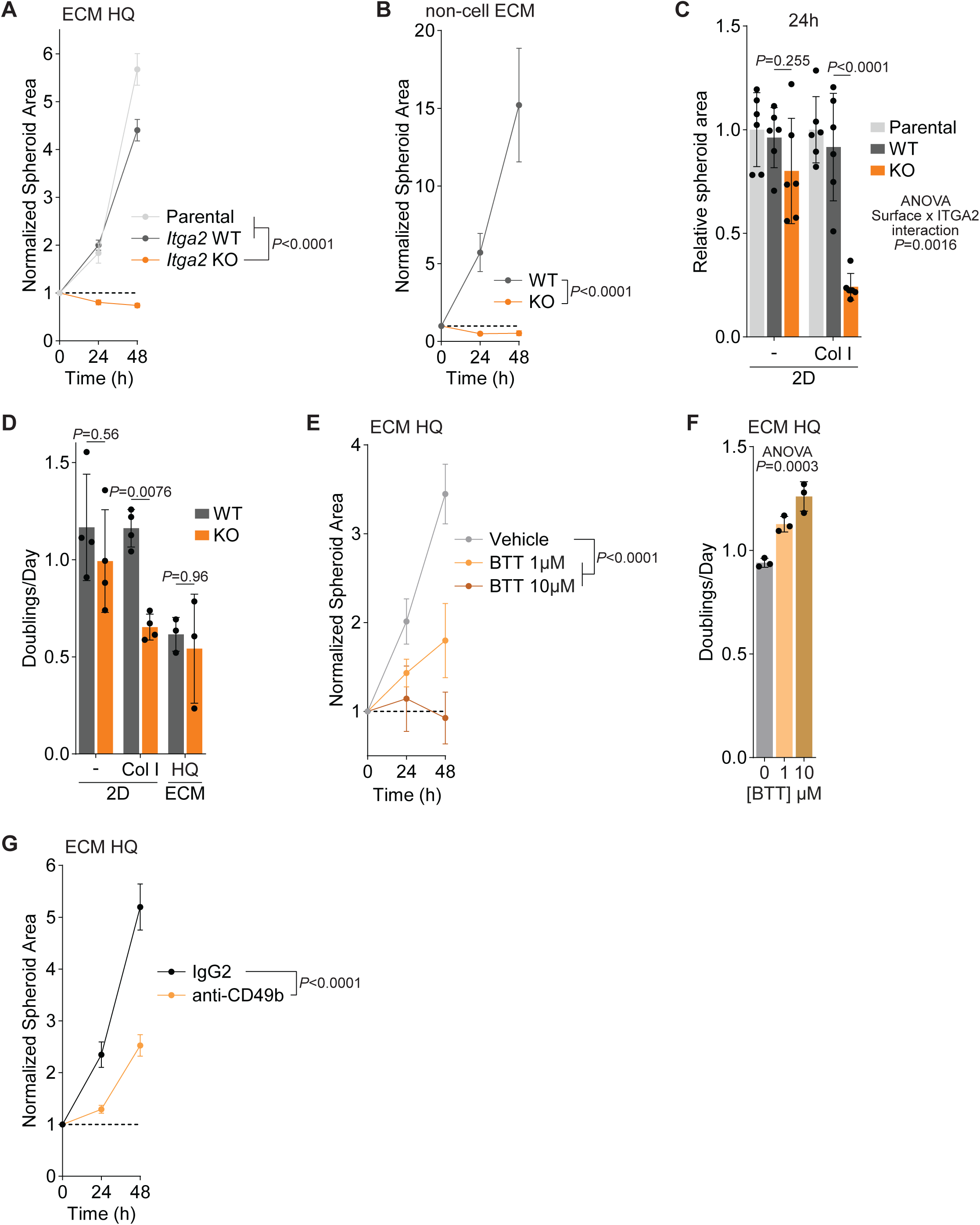
ITGA2 is required for migration but not growth on ECM HQ. (A) Spheroid spreading assay of parental, *Itga2* wild-type (WT) and knockout (KO) KPC2-EYFP spheroids plated on ECM HQ. Spheroid areas normalized to their own initial area after transfer to ECM are shown over 48 h. *n*=6 biological replicates. Data represent mean ± SD. *P* value was calculated by two-way ANOVA. (B) Spheroid spreading assay of *Itga2* WT and KO KPC2-EYFP spheroids on synthetic ECM composed of 4 mg/mL collagen I and 20% Matrigel. Spheroid areas normalized to their own initial area after transfer to ECM are shown over 48 h. *n*=6 biological replicates. Data represent mean ± SD. *P* value was calculated by two-way ANOVA. (C) Spreading assay of PAR, *Itga2* WT and KO KPC2-EYFP spheroids on 2D plastic or collagen I coated surfaces. Normalized spheroid areas after 24 h, relative to PAR spheroids. *n*=6 biological replicates. Data represent mean ± SD. *P* values were calculated by two-way ANOVA and represent the effect of *Itga2* KO on the respective surface, or the overall interaction between the *Itga2* KO and surface effect. (D) Growth of WT and *Itga2* KO KPC2-EYFP cells over 48 h on 2D plastic or collagen I-coated surfaces, and ECM HQ. *n*=4 (2D), *n*=3 (ECM) biological replicates. Data represent mean ± SD. *P* values were calculated by two-way ANOVA (E) Spheroid spreading assay of KPC2-EGFP spheroids plated on ECM HQ in the presence of vehicle or the ITGA2 inhibitor BTT-3033 at 1 or 10 μM. *n*=6 biological replicates. Data represent mean ± SD. *P* value was calculated by two-way ANOVA. (F) Growth of KPC2-EGFP cells on ECM HQ over 48 h in the presence of vehicle or BTT-3033 at 1 or 10 μM. *n*=3 biological replicates. Data represent mean ± SD. *P* value was calculated by one-way ANOVA. (G) Spheroid spreading assay of KPC2-EGFP spheroids plated on ECM HQ in the presence of 20 µg/mL isotype IgG2 or anti-CD49b (ITGA2) blocking antibody. *n*=6 biological replicates. Data represent mean ± SD. *P* value was calculated by two-way ANOVA.

## Discussion

Glutamine availability has been recognized as a key regulator of PDAC cell biology [38]. How it affects the tumor stroma, in particular the ability of fibroblasts to produce ECM, is less well understood. Here, we show that extracellular glutamine regulates the biochemical composition and mechanical properties of fibroblast-derived ECM. ECM made by fibroblasts under glutamine-poor conditions is depleted of fibrillar collagens and collagen maturation enzymes. This ECM is also less stiff than the ECM derived from a glutamine-rich culture. Consequently, changes in glutamine availability impact the ECM’s biological activity, with differential effects on PDAC cell growth and migration. Collectively, these data support a model in which glutamine levels in the tumor microenvironment (TME) can also impact PDAC cell biology indirectly via their effect on CAFs and the ECM they produce. Our findings suggest that PDAC cells may also be able to sense glutamine availability via changes in the ECM, in addition to more direct sensing mechanisms such as GCN2 or mTORC1 [22].

Proteomic analysis of fibroblast-derived ECM indicates a continuum of ECM type and maturation that is coordinated with glutamine availability, from basement membrane towards a more fibrillar and mature ECM (Fig. 1). The enrichment of a basement membrane-like matrisome under LQ conditions, notably collagen IV, may reflect fibroblasts under conditions of cellular stress reverting to a more ancient kind of matrisome [39] essential for cell adhesion and polarity, which are basic requirements for multicellularity. In contrast, when resources (i.e., glutamine) are abundant, fibroblasts may produce the more structurally complex fibrillar ECM required for complex tissue architecture. While we focused our work on PDAC, this shift in ECM composition along the glutamine gradient may originate from fibroblasts’ role in tissue repair. Glutamine levels are reduced in the early phases of wound healing [40]. The coupling of fibrillar collagen synthesis to environmental glutamine may allow inflammation to cease and re-vascularization to occur before new layers of connective tissue are deposited, supporting the efficiency of wound healing.

Our finding that fibrillar collagens are depleted in fibroblast-derived ECM when glutamine is limiting is consistent with prior work demonstrating that TGFβ-induced collagen I synthesis is glutamine dependent [19,20]. We observed that the relative abundance of many matrisome proteins is reduced in low glutamine conditions (Fig. 1D), and this may be explained by lower global translation rates in TGFβ-stimulated fibroblasts in this environment [19]. However, this does not sufficiently explain why fibrillar collagens and associated maturation enzymes are selectively reduced under these conditions, such that, in relative terms, ECM composition is substantially altered in a low glutamine environment. Unlike most other ECM proteins, pro-collagen undergoes extensive post-translational modification (hydroxylation of proline and lysine residues) and processing before it is secreted [41]. If collagen is not hydroxylated correctly, it is degraded in the lysosome [42]. Collagen prolyl- and lysyl-hydroxylation depend on glutamine-derived alpha-ketoglutarate [42], and consistently, alpha-ketoglutarate levels are lower in TGFβ-stimulated fibroblasts under glutamine deprivation [19]. While collagen IV enriched in LQ conditions is also heavily hydroxylated, it has fewer hydroxylation events per chain overall than collagen I with its uninterrupted Gly-X-Y repeats, suggesting a reduced alpha-ketoglutarate demand compared to collagen I.

The selective relative depletion of fibrillar collagens in fibroblast-derived ECM under glutamine limitation (Fig. 1) contrasts with the overall relative accumulation of such collagens in glutamine-poor PDAC. This may be explained by metabolic heterogeneity in PDAC tumors, which display pockets or “nests” of hypoxia and glutamine deprivation [22,43], suggesting that only these areas may show glutamine-driven changes in ECM composition. This idea is supported by the recent identification of spatially confined sub-tumor microenvironments (subTMEs) in PDAC [44], which display altered collagen abundance. Whether glutamine concentrations differ between such subTMEs is unknown. Testing this is challenging given the requirement for advanced tools or analytical techniques, such as regional tumor interstitial fluid sampling or genetic reporters for glutamine stress. Associating collagen levels with glutamine availability in tumors will help clarify whether alterations in the ECM are indeed a consequence of altered perfusion and related changes in tumor nutrient levels, or a cause, as suggested previously [45].

Since the ECM can both promote and restrain tumor progression [46], a key challenge in matrix biology lies in understanding which ECM molecules or properties underlie pro- and anti-tumor effects. Our findings suggest that fibroblast-derived collagen I slows down PDAC cell growth through its crosslinking but promotes PDAC cell migration through its abundance (Fig. 2). The growth-restraining properties of fibroblast-derived ECM in general, and collagen I in particular, are consistent with previous work in pancreatic cancer [15,47]. In addition, it has been demonstrated that the tumor genotype also dictates collagen production and stiffness of fibroblast-derived ECM [48]. This study showed that ECM made by CAFs isolated from p53-mutant tumors, compared to p53-deleted tumors, promotes the invasion of both p53-mutant and p53-deleted cancer cells. The findings that tumor genotype (via changes in paracrine factors) and tumor glutamine levels influence the ECM support the idea that environment-driven changes in fibroblast-derived ECM are sufficient to alter PDAC cell biology.

Here, we describe a glutamine-regulated collagen I-ITGA2-FAK axis that promotes PDAC cell migration. In PDAC, the role of FAK in supporting cancer cell migration is well appreciated [50]. In addition to FAK signaling, other mediators of mechanotransduction, including YAP/TAZ signaling, can transmit signals from the ECM to change cellular behavior [4] and may play a role in the differential effects of ECM HQ and ECM LQ, particularly in response to their altered stiffness. The role of ITGA2 in mediating collagen I-FAK crosstalk is consistent with previous work in ovarian cancer [51]. Consistent with our analysis of TCGA-PAAD data, elevated expression of ITGA2 protein in PDAC patients is associated with poor outcome [52–54]. Interactions between collagen I and ITGA2/ITGB1 promote the growth and migration of PDAC cells [55], consistent with our findings using collagen I as the sole matrix. However, on more physiologically relevant fibroblast-derived ECM, ITGA2 is only required for PDAC cell migration, but not growth, which may be explained by cues from other ECM proteins present. These findings suggest a dominant role of collagen I-ITGA2/ITGB1 interactions in mediating PDAC cell migration on ECM that may be exploited therapeutically.

## Experimental procedures

### Cell Culture

KPC (*Kras*^LSL-G12D/+^;*Trp53*^LSL-R172H/+^;*Pdx1-Cre*) mouse PDAC cells were derived from *KPC* mice and described previously [19,56]. KPC-EYFP and KPC-EGFP are fluorescently labeled derivatives of the parental KPC2 line generated via lentiviral transduction. PSCs were derived from C57BL/6 mice via differential centrifugation and immortalized by spontaneous outgrowth, and were described previously [19,43]. Human PSCs (hPSCs) were obtained from ScienCell (3830) and immortalized by retroviral transduction with the SV40 Large T antigen (Addgene, 13970). 293T cells were obtained from ATCC (CRL-3216). All cells were cultured at 37°C in a humidified incubator with 5% CO_2_ and 20% O_2_. Cells were maintained in high-glucose DMEM (Sigma, D5796) supplemented with 10% FBS (Gemini, 100-106-500), 100 U/mL penicillin, and 100 μg/mL streptomycin (P/S) (Corning, 30002CI). Cells were plated in standard culture media. The following day they were washed with PBS and the media was changed to DMEM without glucose, glutamine and sodium bicarbonate (Sigma, D5030), supplemented with 25 mM glucose (Gibco, A2494001), 3g/L sodium bicarbonate (Fisher, BP328), 10% dialyzed FBS (dFBS) (Gemini, 100-108-500), 1% P/S and the desired concentration of glutamine (Cytiva, SH3003401). For imaging experiments, cells were cultured in FluoroBrite DMEM (Gibco, A1896701) supplemented with 10% FBS, 1% P/S, and 4 mM glutamine. For 2D experiments with collagen I, culture dishes were coated with 0.1 mg/mL rat tail tendon collagen I solution (Corning, 354236). All cell lines were routinely tested for mycoplasma contamination using the MycoAlert Mycoplasma Detection Kit (Lonza, LT07-318) or the MycoStrip Mycoplasma Detection Kit (Invivogen, rep-mysnc-100) and confirmed to be negative. Cells were treated with 1 μM PF-562271 (Sigma, PZ0387), 1 μM PND1186 (Cayman, 17668), 1-10 μM BTT3033 (Tocris, 4724), 0.1 μM Trametinib (Selleck, S2673), 250 μM β-aminopropionitrile (BAPN; Thermo, A1304303), and 20 µg/mL purified Hamster Anti-Rat/Mouse CD49b (BD, 554998).

### Production of non-cell ECM

Non-cell ECMs were prepared by incubating 4 mg/mL or 1 mg/mL high-concentration rat tail collagen I (Corning, 354249) with 2.5% NaOH in PBS and 20% (v/v) GFR Matrigel (Corning, 356231) for 30 minutes at 37°C. The gels were used as a surface for seeding KPC cells or spheroids upon polymerization.

### Production of fibroblast-derived ECM

ECMs were generated as described previously [23]. To enhance matrix adhesion, tissue culture plates were incubated with 0.1% gelatin (Sigma, G1890) for 1 hour at 37°C, followed by a brief PBS wash. Plates were incubated with 1% glutaraldehyde solution (Sigma, G6257) for 30 minutes at room temperature. After three PBS washes, plates were incubated with 1 M ethanolamine (Thermo, 149582500) for 30 minutes at room temperature. Plates were washed thrice with PBS, either used immediately or sealed, and stored in the final PBS wash at 4°C. PSCs were seeded onto coated plates at 8×10^4^ cells/well in 24-well plates, 1.5×10^5^ cells/well in 12-well plates, 4×10^5^ cells/well in 6-well plates, and 8×10^5^ cells per 6 cm dish. Cells were initially seeded in standard DMEM. The following day, the media was changed to DMEM with 10% dFBS and 4 mM (HQ) or 0.4 mM (LQ) glutamine, unless described otherwise, and supplemented with 20 μg/mL sodium ascorbate (Sigma, A4034) and 2 ng/mL TGFβ1 (PeproTech, 100-21). Media were changed every other day for a total of three times. After six days of ECM deposition, plates were decellularized with 0.5% Triton X-100 (Thermo, A16046AP) and 20 mM ammonium hydroxide (Sigma, 221228) in PBS until no intact cells were visible. Plates with ECM were washed four times with PBS and sealed and stored in PBS containing 1% P/S at 4°C until further use. When used for cell seeding, decellularized ECMs were further incubated with 10 μg/mL DNase I (Sigma, DN25) in PBS for 30 minutes at 37°C. ECMs were washed gently three times before cell seeding.

### Analytic Techniques on fibroblast-derived ECM

#### ECM harvest for Western Blot

ECMs were produced and decellularized in 12-well plates as described above. To each well of decellularized ECM, 200 μL preheated 1× LDS sample buffer (Invitrogen, NP0007), supplemented with 1 mM dithiothreitol (DTT), was added. ECMs were scraped off, boiled at 95°C for 10 minutes, and proteins were separated by SDS-PAGE followed by immmoblotting as described below.

#### ECM harvest and preparation for LC-MS

ECMs were produced on 6 cm dishes, decellularized, scraped in PBS, and transferred into low-retention microcentrifuge tubes. After centrifugation and PBS removal, pellets were solubilized in 8 M urea (Sigma, U1250) in 100 mM NH₄HCO₃ (Thermo, 393212500) and reduced with 10 mM DTT (Thermo, A39255) at 37°C for 2 hours at 1400 rpm on a shaker. Samples were cooled to room temperature, alkylated with 25 mM iodoacetamide (Thermo, A39271) for 30 minutes in the dark at room temperature, then diluted to 2 M urea in 100 mM NH₄HCO₃. 2000 U//100 µL PNGase F (New England Biolabs, P0704S) was added and incubated for 2 hours at 37°C, followed by addition of 2 µg/100 µL Lys-C (Thermo, 90307) with incubation for 2 hours at 37°C, and 6 µg/100 µL trypsin (Thermo, 90058) with incubation overnight at 37 °C and 1400 rpm, shaking. Trypsin was added a second time at 3 µg/100 µL with incubation for 2 h at 37°C to improve the digestion of cross-linked ECM. Digestions were quenched by acidification with 50% trifluoroacetic acid (Thermo, 85183) to pH<2 and centrifuged at 16,000g for 5 minutes at room temperature. Clarified peptides were desalted using Pierce™ Peptide Desalting Spin Columns (Thermo, 89851), dried in a speedvac vacuum concentrator (Thermo, SPD140P2115), and reconstituted in 0.1% formic acid (Thermo, 28905) and 5% acetonitrile (Avantor, BDH83639.400) in water. Peptide concentration was measured with the Pierce™ Quantitative Colorimetric Peptide Assay (Thermo, 23275).

#### Liquid chromatography Mass-Spectrometry (LC-MS) proteomics

LC-MS-based matrisome proteomics was conducted by the Mass Spectrometry Core Facility at the University of Illinois as previously described [57]. Peptide samples were analyzed using a Q Exactive HF mass spectrometer coupled with an UltiMate 3000 RSLC nanosystem with a Nanospray Frex Ion Source (Thermo). Digested peptides were loaded into a Waters nanoEase M/Z C18 (100Å, 5um, 180um x 20mm) trap column and then a 75 μm x 150mm Waters BEH C18 (130A, 1.7um, 75um x 15cm) and separated at a flow rate of 300nL/min. Solvent A was 0.1% FA in water, and solvent B was 0.1% FA, 80% ACN in water. The solvent gradient of LC was 5% B in 0-3 min, 8% B in 3.2min, 8-40% B in 110 min, 40-95% B in 119min, wash 95% in 129 min, followed by 5% B equilibration until 140 min.

Full MS scans were acquired in the Q-Exactive mass spectrometer over a 350-1400 m/z range with a resolution of 120,000 (at 200 m/z) from 10 min to 120 min. The AGC target value was 3.00E+06 for the full scan. The 20 most intense peaks with charge states 2, 3, 4, 5 were fragmented in the HCD collision cell with a normalized collision energy of 28%, and these peaks were then excluded for 30s within a mass window of 1.2 m/z. A tandem mass spectrum was acquired in the mass analyzer with a resolution of 30,000. The AGC target value was 1.00E+05. The ion selection threshold was 2.00E+04 counts, and the maximum allowed ion injection time was 50 ms for full scans and 120 ms for fragment ion scans.

Spectra were searched against the mouse database using MaxQuant (2.0.3.1) with the following parameters: parent mass tolerance of 10 ppm, constant modification on cysteine alkylation, hydroxylation of proline and lysine, cyclization of Gln to pyroGlu, variable modification on methionine oxidation, deamidation of asparagine and glutamine. Search results were entered into Scaffold DDA software (v6.6.0, Proteome Software, Portland, OR) for compilation, normalization, and comparison of spectral counts, etc. The filtering criteria of protein identification were a 1% false discovery rate (FDR) of protein and a minimum peptide count of one.

#### Atomic Force Microscopy (AFM)

Fibroblast-derived ECMs were produced on glass-bottom 6 cm dishes (Thermo, 150682) and decellularized. AFM was conducted on decellularized ECMs in PBS by the Molecular Cytology Core Facility at Memorial Sloan Kettering Cancer Center as previously described [58]. Experiments were performed with an MFP-3D-BIO AFM (Oxford Instruments), integrated with an inverted Zeiss AxioObserver Z1 microscope and an AxioCam. The Zeiss microscope was used to position the AFM cantilever visually with respect to the sample. AFM scanning was performed on several random areas across five independent ECMs per glutamine concentration. Before each experiment, the cantilever spring constant was calibrated using the thermal noise method, and the optical sensitivity was determined using a glass-bottom Petri dish filled with PBS as an infinitely stiff substrate.

### Characterization of Cancer Cell Behavior on ECM

#### Seeding Cells on ECM

To assess the proliferative behavior of cancer cells on ECMs generated under different glutamine concentrations, KPC-EYFP cells were seeded at 1×10^4^ cells/well onto decellularized ECMs in 24-well plates. Drug treatments were applied the day after seeding. To quantify growth, cell numbers were counted at this point to establish a baseline, followed by a second count 48 hours later. ECMs were enzymatically digested in all conditions (including plastic controls) using 40 µg/mL collagenase P (Sigma, 11249002001) in 0.25% Trypsin-EDTA (Sigma, T4049). Total viable cell numbers were measured using a CASY cell counter (OMNI Life Science) with manual gating to exclude debris and dead cells, and doubling time was calculated using the following formula:

Doubling time (in days) = (t2 - t1) × log(2) / [log(N2) - log(N1)]

where *N1* and *N2* are the viable cell counts on the day after seeding and the day of the final count, respectively, and *t2 - t1* is the time interval in days.

#### Spheroid spreading assay

KPC-EGFP or KPC-EYFP cells were seeded into ultra-low attachment 96-well plates (Corning, 4515) at 1.2×10^4^ cells/well to allow for spheroid formation overnight. Intact spheroids were transferred at one spheroid/well onto decellularized ECM, non-cell ECM, or collagen I-coated 24-well plates using a wide-bore P1000 pipette tip. Immediately after transfer, spheroids were examined for integrity by phase-contrast microscopy, and only intact spheroids were retained for further analysis. Drug treatments were administered at the time of transfer. All spheroid cultures were conducted in FluoroBrite DMEM to improve fluorescence imaging quality and minimize background signals. Spheroids were imaged 4h after seeding (0h time point in graphs) and 24 hours and 48 hours later using a Keyence BZ-X800 fluorescence microscope with a 4x objective under oblique illumination. Both brightfield and GFP fluorescence channels were captured for each well. Exposure and brightness were auto-calibrated, and the fluorescence signal was verified to align with the brightfield image.

Spheroid images were analyzed using a custom batch-processing macro written in ImageJ Macro Language (IJM). Each image was first rescaled by setting the spatial calibration to 121 pixels per 500 μm. Images were converted to 8-bit grayscale, and contrast was normalized to minimize brightness variation. A Gaussian blur (σ = 2) was applied to reduce high-frequency noise, followed by background subtraction using a sliding paraboloid with a 500-pixel rolling radius. Spheroids were segmented using automatic thresholding with the Otsu method, and the resulting binary masks were refined by filling holes and applying a single round of dilation to ensure continuity of the segmented regions. Spheroid areas were identified using the *Analyze Particles* function in ImageJ, with a minimum area cutoff of 500 pixels to exclude noise. The macro computed the total area for each well by summing individual spheroid regions and converting the values from pixels to square micrometers using the defined spatial calibration. Results for each image were appended to a CSV file that recorded the experimental condition, well position, and total spheroid area. The compiled dataset was then used for downstream statistical analysis and visualization.

#### Western Blot

Cells cultured on decellularized ECM were harvested in 1× RIPA buffer (Sigma, 20-188) supplemented with 1× protease and phosphatase inhibitors (Thermo, 78447). After 15 minutes of incubation on ice, cells were scrapped, transferred into microcentrifuge tubes, and lysates were cleared by centrifugation at 18,000 × g for 10 minutes at 4°C. In some experiments, subcellular fractionation was performed. Cells on ECM were scrapped with ice-cold PBS into microcentrifuge tubes, pelleted at 1,000 × g for 3 minutes, and resuspended in ice-cold Wash Buffer (10 mM HEPES-KOH (pH 7.9), 0.5 mM KCl, 0.1 mM EDTA, and 0.1 mM EGTA) supplemented with 1:100 NP-40 and 1× protease and phosphatase inhibitors. Cell suspensions were incubated on ice for 5 minutes, followed by a 5-second vortex to disrupt membranes. Samples were centrifuged at 1,000 × g for 5 minutes to pellet nuclei. A 135 μL aliquot of the resulting supernatant was transferred to new tubes and designated as the cytosolic fraction. The nuclear pellet was washed once with Wash Buffer lacking NP-40, centrifuged again at 1,000 × g for 5 minutes, and the supernatant was discarded. The pellet was resuspended in 50 μL Nuclear Lysis Buffer (10 mM HEPES-KOH (pH 7.9), 10 mM NaCl, 0.1 mM EDTA, 0.1 mM EGTA, 0.1% NP-40) containing 1× protease and phosphatase inhibitors, 0.1% SDS, and 0.5 μL of benzonase (Sigma, 70746). After 15 minutes incubation on ice, both cytosolic and nuclear fractions were cleared by centrifugation at 15,000 × g for 5 minutes. Protein concentrations were quantified by BCA assay (Thermo, 23227) prior to SDS-PAGE and immunoblotting.

The following primary antibodies were used: FAK p-T397, FAK (1:1,000; Cell Signaling, 13009), ERK1/2 p-T202/Y204 (1:1,000; Cell Signaling, 4377), ERK1/2 (1:1,000; Cell Signaling, 9107), mouse Collagen I (1:1,000; Abcam, ab21286), human Collagen I (1:1,000; Proteintech, 67288-1-Ig), Fibronectin (1:1,000; Abcam, ab2413), ITGA2 (1;1;000; Abcam, ab181548), β-Actin (1:5,000; Sigma, A5441), Vinculin (1:5,000; Sigma, V9131). The following secondary antibodies were used: anti-rabbit IgG, peroxidase-linked (1:5,000; Cytiva, NA9341ML), and anti-mouse IgG, peroxidase-linked (1:5,000; Cytiva, NA9311ML).

### Genetic manipulation of ITGA2 expression

#### Viral infection

Lentiviral particles were produced in 293T cells by using psPAX2 (Addgene, 12260) and pCMV-VSV-G packaging plasmids (Addgene, 8454). Retroviral particles were produced in 293T cells by using pCG-gag-pol (Addgene, 14887) and pCMV-VSV-G packaging plasmids. Media was changed the next day, and viral supernatant was collected 48 hours later, passed through a 0.45 μm nylon filter and used for transduction in the presence of 8 μg/ml polybrene (Sigma, 107689).

#### CRISPR-Cas9-mediated deletion of Itga2

Single-guide RNAs (sgRNAs) targeting *Itga2* were designed using GuideScan (http://www.guidescan.com/) and cloned into the lentiCRISPR v2 vector (Addgene, 52961). The following target sequence was used: TCTGAGACGCGCCAACATGG. KPC-EGFP cells were transduced as described above and subjected to 2 μg/ml puromycin (Sigma, P4512) antibiotic selection two days later. To isolate a clonal *Itga2* knockout line, cells from the selected pool were seeded at approximately one cell per well into 96-well plates and expanded into larger wells. Clonal isolates were screened by Western blot to confirm loss of ITGA2 protein expression. Two independent pairs of *Itga2* WT and KO clones were used for experiments, with representative data from one pair shown.

#### Expression of human ITGA2

Human *ITGA2* (*hITGA2*) cDNA containing plasmid was obtained from Sino Biological (HG13024-M) and subcloned into a modified version of pTRE-Tight (Takara, 631059) with addition of a C-terminal FLAG tag using Gibson assembly. KPC2-EYFP *Itga2*-KO cells were transduced as described above and subjected to 500 μg/ml hygromycin B (Invivogen, ant-hg-1) antibiotic selection two days later. Inducible expression of *hITGA2* was achieved with 1 µg/mL doxycycline (Fisher, BP26531) treatment, started one passage before cells were plated for experiments and maintained throughout.

### TCGA analysis

TCGA PAAD gene-level expression data (HiSeqV2) were obtained from the UCSC Xena repository, and the mean expression of *COL1A1* and *COL1A2* was calculated for each tumor. Samples were stratified into *COL1A1/2*^high^ and *COL1A1/2*^low^ groups using the median expression as a cutoff. Differential expression analysis between the two groups was performed using the *limma* package in R 4.4.1. Genes were ranked by log2-fold change and subjected to gene set enrichment analysis (GSEA) with *clusterProfiler* using curated pathway collections from MSigDB. Enrichment was specifically assessed for signaling pathways implicated in extracellular matrix– integrin signaling. The correlation between *COL1A1/2* and *ITGA2* expression was evaluated using Pearson correlation and visualized by scatterplot regression. Survival analysis was performed with Survival Genie 2.0, using the optimal cut-off method.

### Single-cell RNA Sequencing (scRNA-seq) Analysis

scRNA-seq data from human PDAC were previously collected by Peng et al. [30] and is accessible from the Genome Sequence Archive under project PRJCA001063. Quality control, merging, doublet removal, and cell annotations were previously performed by Khaliq et al. and are available at https://zenodo.org/records/10712047, under the filename ‘scRNA-seq_Data_post_qc.rds’ [59]. Downstream analysis was performed using R 4.4.1 and the following R packages: “ggpubr_0.6.1”, “RColorBrewer_1.1-3”, “dplyr_1.1.4”, “ggplot2_3.5.2”, “Seurat_5.3.0”, “SeuratObject_5.1.0”, “sp_2.2-0”, “readr_2.1.5”, “CellChat_1.6.1”, “ComplexHeatmap_2.22.0”, “patchwork_1.3.1”, “Biobase_2.66.0”, “igraph_2.1.4”, and “BiocGenerics_0.52.0”.

PDAC cells were subsetted using *Seurat’s subset()* function based on tumor tissue status, stored in metadata as “orig.ident”. Cell types representing less than 0.05% of the remaining population were excluded. Fibroblasts were similarly subsetted based on cell type annotation, stored in metadata as “cell_type_new”. Following each subset, remaining cells were processed using the standard Seurat workflow wherein genes were scaled and centered using *ScaleData(),* highly variable features were found using *FindVariableFeatures(),* and principal component analysis (PCA) was performed using *RunPCA().* Cell clusters were calculated and identified using *FindNeighbors()* and *FindClusters()* with parameters determined by ElbowPlot(). RunUMAP() was used to generate the UMAP, and *DimPlot()* was used to visualize the result. For single gene feature expression, *FeaturePlot()* was used for UMAP, and *VlnPlot()* was used for expression probability distribution visualizations. For signature scores, *AddModuleScore()* was used for calculations, and results were visualized in the same way as single gene features, but with limits on expression to exclude outliers (>99.95% or <0.05% signature score) [60].

To infer cell communication networks, including ligand–receptor interactions, the standard CellChat workflow and parameters were used [61]. Differential interaction estimates were calculated with the entire CellChatDB using *identifyOverExpressedInteractions().* Communication probability and cellular communication networks/pathways were estimated with *computeCommunProb()*, *filterCommunication()*, and *computeCommunProbPathway()*. Communication networks between fibroblasts and other cell types were visualized as an aggregated cell-cell communication network using *netVisual_circle()*, displaying weight. Global collagen signaling networks between ligands and receptors of different cell types were visualized using *netVisual_heatmap()* and *netVisual_bubble().* Specific collagen communication network Ligand-Receptor pairs were plotted using *netVisual_chord_gene()*.

### Statistical Analysis

An unpaired, two-tailed *t-test* was applied to compare one variable between two groups. One-way ANOVA was applied to compare one variable between three or more groups. Two-way ANOVA was applied to compare two variables between two groups. Correction for multiple comparisons was done using the Dunnett or Sidak method, as appropriate. Statistical analysis was done in GraphPad Prism 10 or R 4.4.1. Data with *P*<0.05 was considered statistically significant. Graphs show the mean ± SD with individual datapoints, unless indicated otherwise in the figure legends. Unequal sample size between conditions results from different starting sample numbers or differences in available ECMs. ECMs that were damaged during production were not used for subsequent experiments.

## Acknowledgements

We thank all members of the Schwӧrer laboratory for helpful discussions. We thank Dr. Alexandra Naba and James Considine for advice on sample preparation for matrisome proteomics. We thank Drs. Hui Chen and Lasanthi Jayathilaka from the Mass Spectrometry Core Facility at the University of Illinois, Chicago (UIC), for proteomics services, and Dr. Katia Manova and Biran Wang from the Molecular Cytology Core Facility at Memorial Sloan Kettering Cancer (MSKCC) for atomic force microscopy services.

## Author contributions

K.T.: Investigation, Methodology, Formal Analysis, Visualization, Writing

J.G.: Investigation, Methodology, Formal Analysis, Writing

M.R.S.D.: Formal Analysis, Methodology, Visualization, Writing

S.L.: Investigation, Formal Analysis

Y.C.H.: Investigation

K.M.F.: Methodology

S.S.: Conceptualization, Investigation, Methodology, Resources, Formal Analysis, Visualization, Writing, Supervision, Funding Acquisition, Project Administration

## Funding

The UIC Research Resources Center – Mass Spectrometry Core Facility was established in part by a grant from The Searle Funds at the Chicago Community Trust to the Chicago Biomedical Consortium and support from an NIH S10 shared instrumentation grant (1S10OD027016-01). The Molecular Cytology Core Facility at MSKCC is supported by the Cancer Center Support Grant from the NCI (P30CA008748). This work was supported by grants from the NCI (R00CA259224, to S.S.; T32CA009594, to M.R.S.D.), the Department of Defense (HT94252410401, to S.S.), the AGA Research Foundation (Augustyn Award in Digestive Cancer, to S.S.), the Cancer Research Foundation (to S.S.), and the NIGMS (K12GM146658, to K.M.F.). S.S. is also supported by the University of Chicago Medicine through the Beverly Duchossois Cancer Fund. This work was partly supported by the University of Chicago-Taiwan Science (UCTS, to Y.C.H.) Program, made possible through the generosity of our donors.

## Conflict of interest

The authors declare no competing interest.

---

**Supplementary Figure 1:**
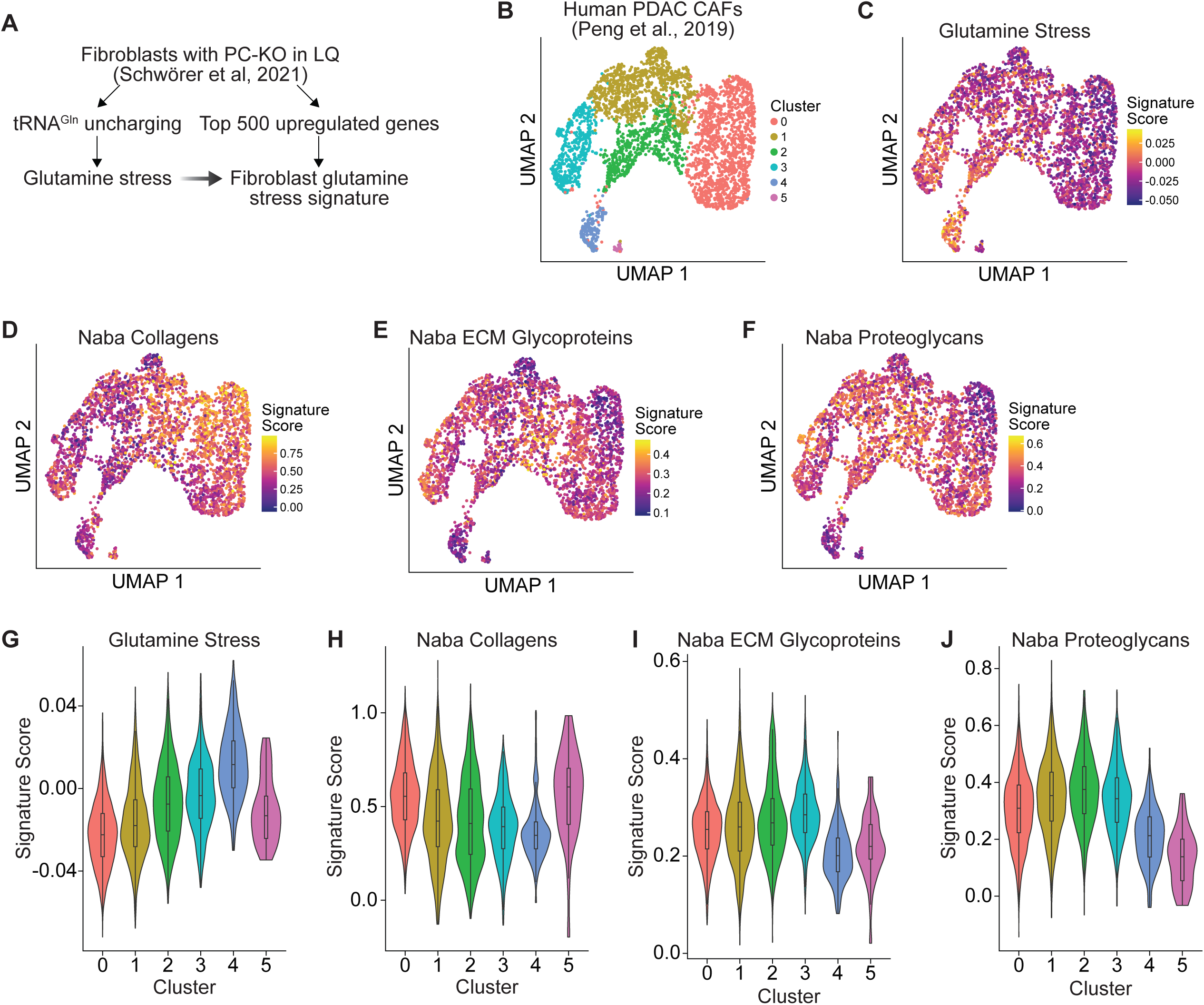
Glutamine availability is associated with reduced collagen expression in CAFs from PDAC patients. (A) Derivation of a fibroblast glutamine-stress gene signature. The top 500 (log2 fold change) upregulated genes in *PC* KO compared to control NIH-3T3 cells cultured in 0.2 mM glutamine (GSE169588) were defined as the “Fibroblast glutamine-stress” signature, given that PC KO cells display tRNA^Gln^ uncharging under this condition. (B) UMAP of human PDAC CAFs subsetted from the sc-RNA-seq dataset (PRJCA001063) showing six CAF clusters (0–5). The complete dataset is shown in Sup. Fig. 5A. (C–F) Signature scores projected on the CAF UMAP from (B): (C) glutamine-stress signature obtained from (A), (D-F) matrisome gene sets from the matrisome database. (G-J) Violin plots showing per-cell score distributions of signatures in (C-F) across CAF clusters 0 to 5.

**Supplementary Figure 2.**
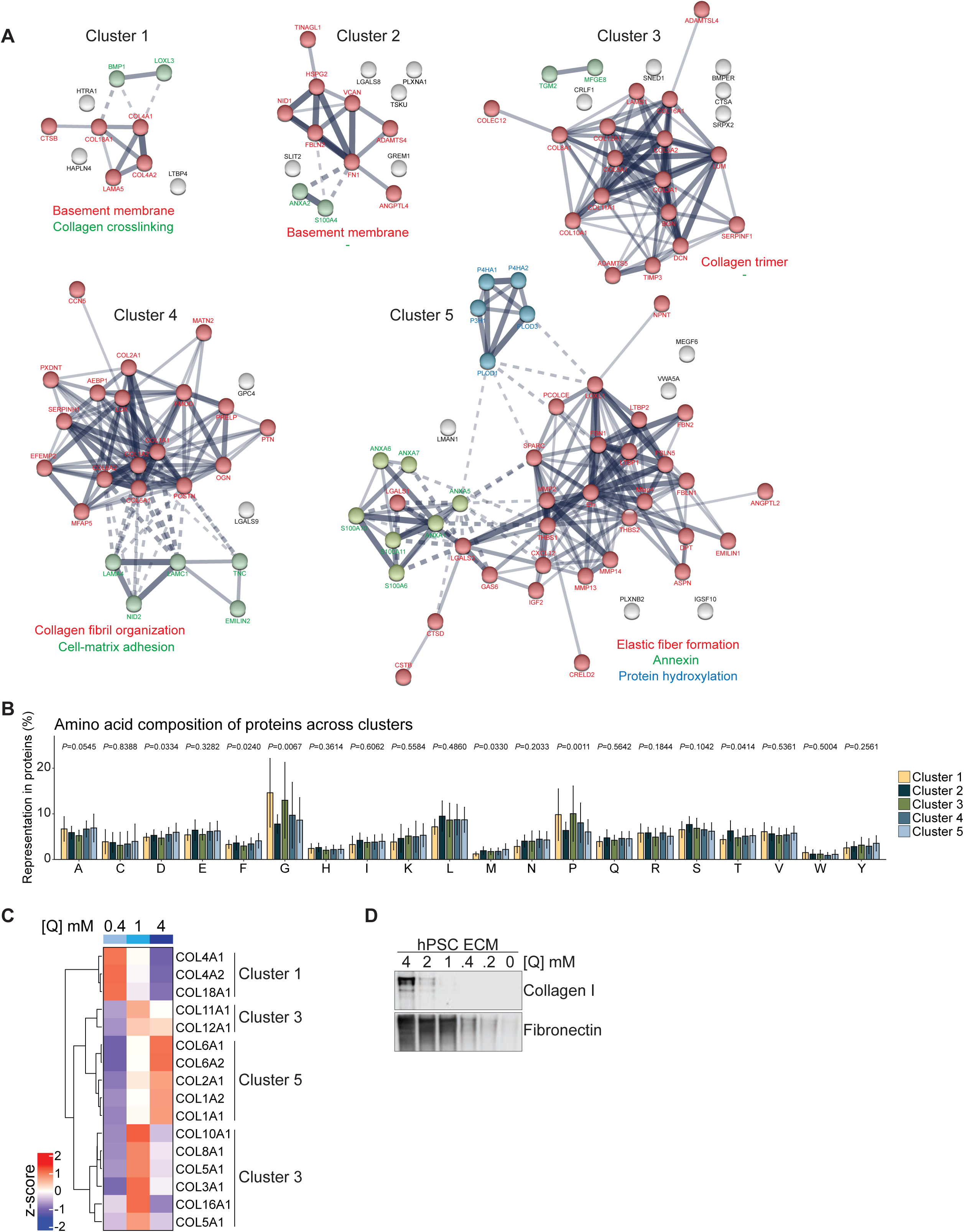
(related to Figure 1): ECM proteomics. (A) Protein-protein interaction networks (STRING) for the five clusters identified in Fig. 1D, with selected functional annotations of sub-clusters indicated with matching color. “-“ indicates the absence of a common annotation for the sub-cluster members. Edge thickness denotes interaction confidence. (B) Amino acid composition of proteins across clusters. Bars show the % representation of each amino acid within cluster proteins. *P* values for each amino acid were calculated by two-way ANOVA. (C) Heatmap showing z-score average precursor intensities for collagens across 4-, 1-, and 0.4 mM glutamine (Q), with cluster assignment from Fig. 1D indicated. Order follows Euclidean distance. *n*=3 biological replicates. (D) Western blot of human PSC (hPSC)-derived ECM across the indicated glutamine concentrations. ECMs were collected in the same volume, and equal volumes unadjusted for protein concentrations were loaded. Representative image.

**Supplementary Figure 3.**
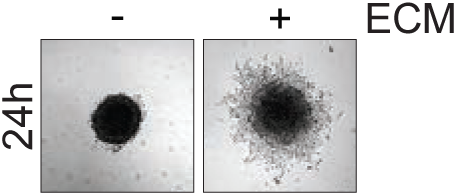
(related to Figure 2): Spheroid spreading assay. Representative brightfield images of KPC2-EGFP spheroids cultured on top of ECM HQ or plastic for 24 h.

**Supplementary Figure 4.**
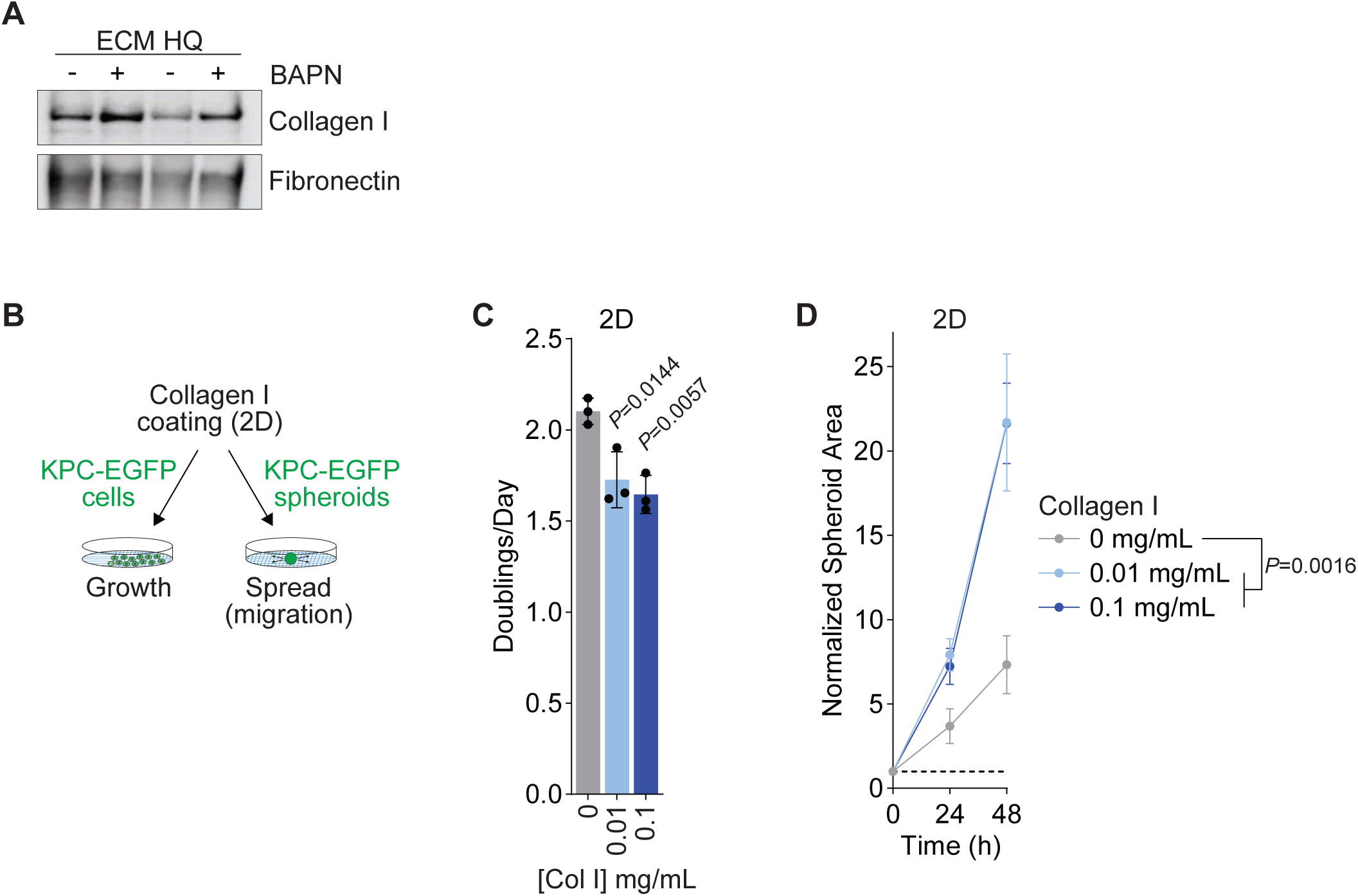
(related to Figure 3): Collagen I is sufficient to promote PDAC cell migration. (A) Western blot of ECM HQ prepared ± 250 µM β-aminopropionitrile (BAPN). Representative image. (B) Schematic of 2D assays using collagen I-coated plates. (C) Growth of KPC2-EGFP cells on plates coated with the indicated collagen I concentrations. Bars show mean ± SD. *n*=3 biological replicates. *P* values were calculated by one-way ANOVA. (D) Spreading of KPC2-EGFP spheroids over 48 h on 2D plates coated with the indicated collagen I concentrations. Spheroid areas normalized to their own initial area after transfer to ECM. *n*=3 biological replicates. Data represent mean ± SD. *P* value was calculated by two-way ANOVA.

**Supplementary Figure 5.**
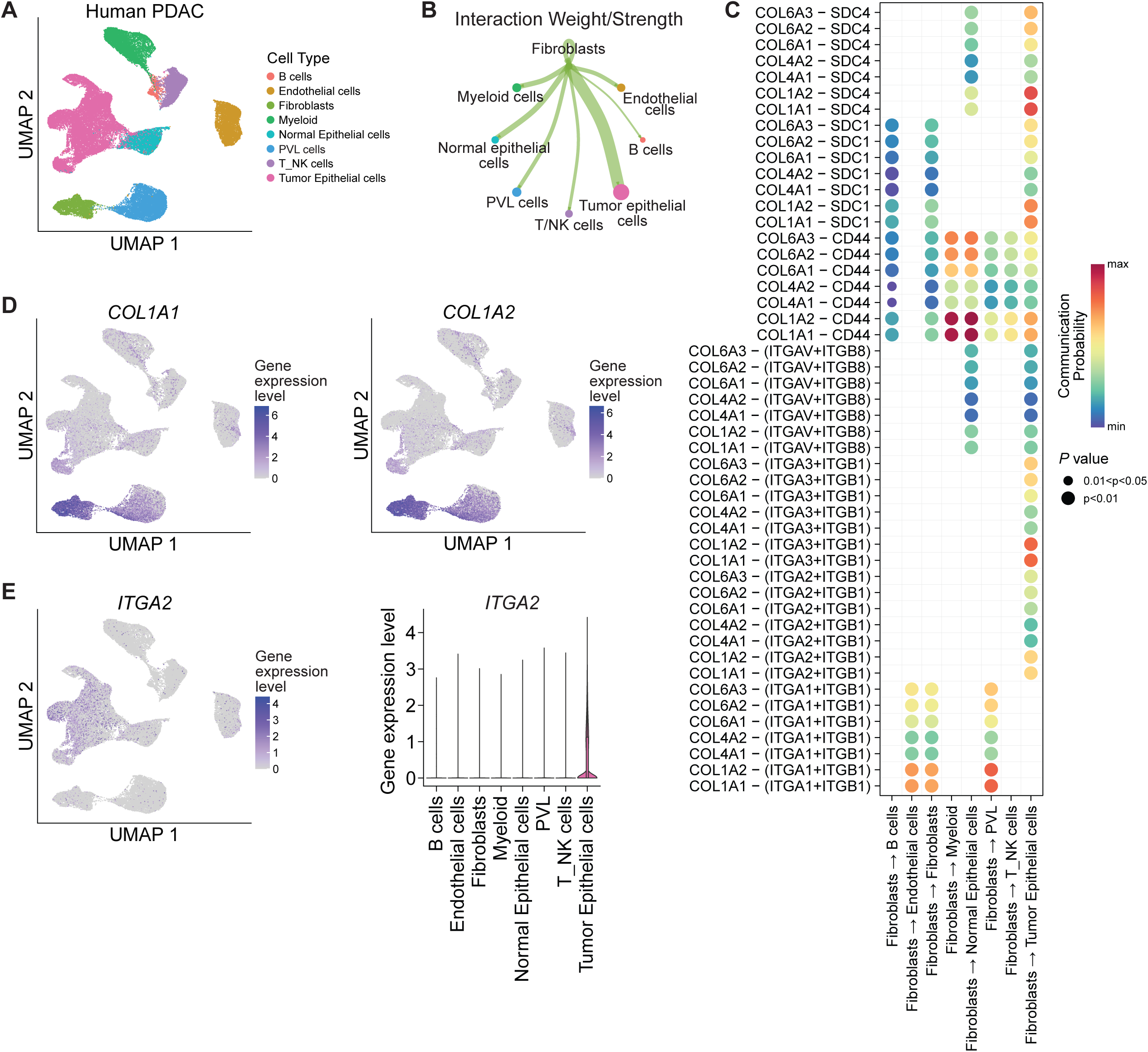
(related to Figure 5): Cell chat analysis of human PDAC to infer fibroblast interactions. (A) UMAP of the human PDAC dataset (PRJCA001063) showing major cell types. (B) Outgoing interaction strength from fibroblasts to the indicated cell types. (C) Bubble plot of predicted collagen ligand-receptor pairs from fibroblasts to the indicated recipient cell types. (D) UMAP feature plots for *COL1A1* and *COL1A2* expression. (E) UMAP feature plot and cell-type violin plot for *ITGA2* expression.

**Supplementary Figure 6.**
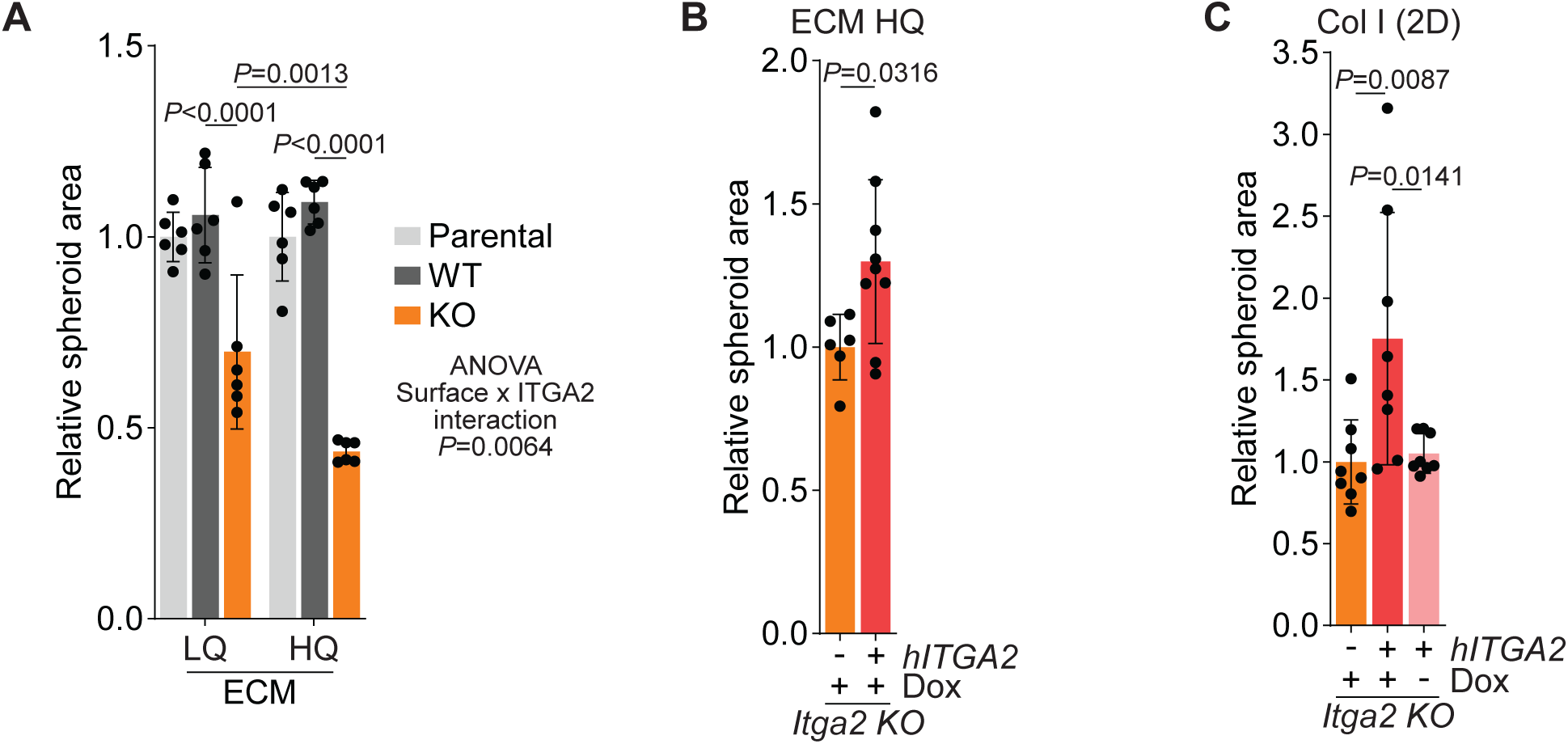
(related to Figure 6): Controls for *Itga2* KO effects on PDAC cell migration. (A) Spreading assay of parental, *Itga2* wild-type (WT) and knockout (KO) KPC2-EYFP spheroids on ECM LQ and ECM HQ. Normalized spheroid areas after 24 h, relative to parental spheroids. *n*=6 biological replicates. Data represent mean ± SD. *P* values were calculated by two-way ANOVA and represent the effect of *Itga2* KO on the respective surface, or the overall interaction between the *Itga2* KO and surface effect. (B) Spreading assay of *Itga2* KO KPC2-EYFP spheroids expressing empty vector or DOX-inducible *hITGA2*, on ECM HQ in the presence of 1 µg/mL doxycycline. Normalized spheroid areas after 24 h, relative to empty vector. *n*=6 (empty vector), *n*=9 (hITGA2). Data represent mean ± SD. *P* value was calculated by two-tailed, unpaired *t* test. (C) Spreading assay of *Itga2* KO KPC2-EYFP spheroids expressing empty vector or DOX-inducible *hITGA2*, on collagen I-coated plates in the presence or absence of 1 µg/mL doxycycline. Normalized spheroid areas after 24 h, relative to empty vector. *n*=8 biological replicates. Data represent mean ± SD; *P* values were calculated by one-way ANOVA.

## Notes

### Competing Interest Statement

The authors have declared no competing interest.

